# Visual recognition of the female body axis drives spatial elements of male courtship in *Drosophila melanogaster*

**DOI:** 10.1101/576322

**Authors:** Ross M McKinney, Yehuda Ben-Shahar

## Abstract

Like other mating behaviors, the courtship ritual exhibited by male *Drosophila* towards a virgin female is comprised of spatiotemporal sequences of innate behavioral elements. Yet, the specific stimuli and neural circuits that determine when and where males release individual courtship elements are not well understood. Here, we investigated the role of visual object recognition in the release of specific behavioral elements during bouts of male courtship. By using a computer vision and machine learning based approach for high-resolution analyses of the male courtship ritual, we show that the release of distinct behavioral elements occur at stereotyped locations around the female and depends on the ability of males to recognize visual landmarks present on the female. Specifically, we show that independent of female motion, males utilize unique populations of visual projection neurons to recognize the eyes of a target female, which is essential for the release of courtship behaviors at their appropriate spatial locations. Together, these results provide a mechanistic explanation for how relatively simple visual cues could play a role in driving both spatially- and temporally-complex social interactions.

## Introduction

Courtship and other social interactions between conspecifics often depend on ritualistic spatio-temporal transitions between distinct innate behavioral elements [1, 2]. Yet, the specific sensory stimuli and neuronal circuits that drive the spatial and temporal aspects of social interactions remain unknown for most species. During courtship behaviors, many animals rely on the visual system to identify salient patterns, colors, or motion cues from conspecifics that promote or inhibit mating-related behaviors [3, 4]. In diverse insect species, motion cues have been shown to be particularly important for triggering male chase behaviors during courtship [5, 6, 7, 8, 9]. These motion cues are detected and processed by visual projection neurons in the brain, which connect to downstream motor centers to generate relevant behavioral outputs [10, 11, 12]. While motion detection is important for keeping the male in proximity to a moving target during courtship, how specific visual cues and neural pathways might regulate the proper spatio-temporal coordination of other mating displays remains largely unknown.

In the fruit fly *Drosophila melanogaster*, the copulation success of males depends on a premating courtship ritual that consists of a sequence of stereotyped behavioral elements including chasing, orienting, singing, scissoring, tapping, licking, and attempted copulation [13, 14, 15]. Although different courtship elements are somewhat independent of one another, there are strong temporal inter-relationships between individual behavioral elements, where the probability of transitioning from one behavior to another is relatively fixed [1]. However, sensory deficits can affect the frequency of these transitions and can also impact both courtship latency and intensity [16, 17, 2]. Similarly, male mating behaviors include important spatial components which support successful copulation, including extension of male’s wing nearest the female’s body to allow for improved perception of auditory cues by the female [17, 18, 9]. However, the specific sensory modalities and cues that drive the appropriate spatial and temporal patterns of individual courtship elements, as well as the transitions between specific elements, are not well understood.

Previous work has suggested that visual recognition of female motion is sufficient to trigger male courtship and that males use motion cues to orient towards and chase target females [6, 7, 8, 9, 12]. Recently, a subset of visual projection neurons in the Lobula Column (LC) of the male brain was shown to be tuned to female-like movements and specifically required for the proper orientation of a male towards a female during courtship [12]. However, which and how other motion-independent visual cues contribute to male courtship remains mostly unknown.

Here we investigated the visual features and neural circuits that regulate both spatial and temporal components of the courtship ritual in *Drosophila* males. By using computer vision- and machine learning-based analyses of male courtship behaviors towards stationary female targets, we demonstrate that the timing and positioning of males during specific courtship elements depends on visual signal processing. Specifically, we show that males use the eyes of their courtship targets as a visual guide to release bouts of tapping, scissoring, and orienting at appropriate times and spatial locations surrounding the female. Further, we find that the spatial positioning of the male depends on the activity of several classes of LC neurons in the visual system. Together, these data suggest that *Drosophila* males use visual recognition of morphological features present on the female’s body to regulate appropriate spatiotemporal distributions of particular courtship behavioral elements.

## Results

### Vision is required for stereotyped behavioral positioning during courtship

Female motion is an important visual cue which male flies use to initiate and direct chase behaviors during bouts of courtship [6, 8, 9, 12]. However, whether other visual cues also play a role in regulating spatial or temporal aspects of courtship elements is mostly unknown. Therefore, here we hypothesized that in addition to responding to motion, males are capable of visually recognizing anatomical features present on the female to direct courtship elements to distinct spatial locations. To separate the effects of female body morphology from motion on the spatial positioning of the male during courtship, we developed a simplified courtship paradigm which eliminates female motion-related visual cues and used custom tracking and behavioral classification software to analyze the spatial localization of males during specific elements of the courtship ritual (Figure 1A). We further used this software to analyze the temporal inter-relationships between specific courtship behaviors (Supplemental Figure 1).

**Figure 1:**
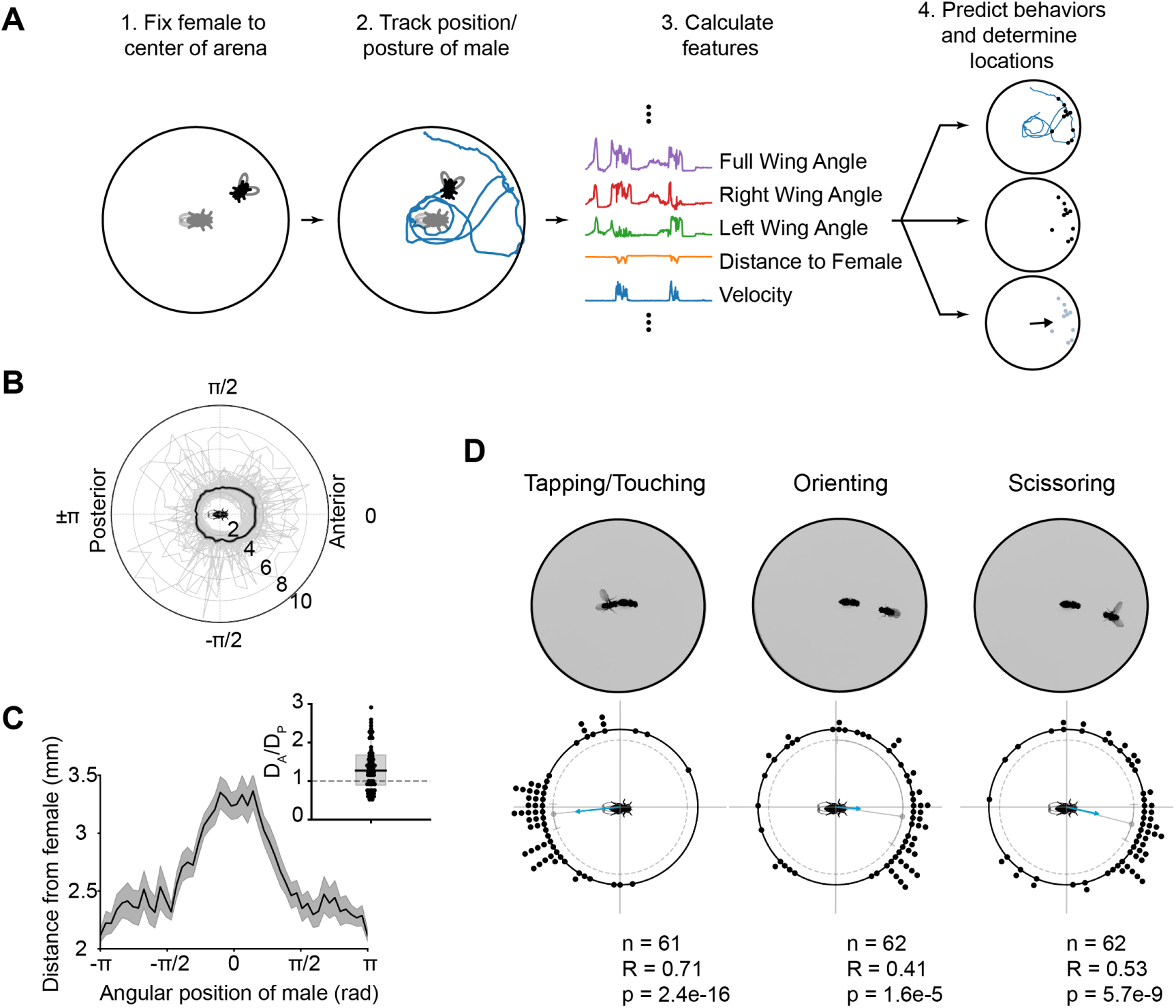
Male courtship behaviors occur at stereotyped locations around the female. **(A)** Overview of algorithm used to determine locations of male mating behaviors during courtship. (1) A female is first fixed to the center of a courtship arena, and then a male is introduced and allowed to court the female for 10 minutes. (2) The male is tracked, and (3) the tracked features are used in a boosted decision tree classifier to predict frames containing a behavior of interest. (4) The position of the male in positively-classified frames is determined, and the mean behavioral position over the trial is recorded. **(B)** The average courtship path of Canton-S males (n=62) over the course of a courtship trial. Each thin gray line represents the average path of an individual male. The thick black line represents the mean of all males. Note that for each behavioral trial, all male tracks have been rotated with respect to the female such that the females anterior-posterior axis is aligned along the horizontal with the anterior end near 0 and the posterior end near *θ* rad. **(C)** The mean courtship path (same as in B) of Canton-S males, shown in Cartesian coordinates. **(C, inset)** The ratio of the maximum male-female distance when the male is on the anterior half of the female (*D*_*A*_) to when the male is on the posterior half of the female (*D*_*P*_). Note that this is significantly greater than 1 (*p* < 0.001, 1-Sample T-Test). **(D)** Examples of individual frames containing positively-classified behaviors are shown above mean angular locations, across flies, for each behavior. Each black point represents the mean behavioral location of an individual fly, and the direction of the arrow represents the mean behavioral location of all flies. The length of the arrow is proportional to the Rayleigh R-value for the total population of flies. The median and 95% confidence intervals surrounding the median are shown as a gray point and lines just beneath the points representing the individual flies.

Using this assay, we first characterized the spatial aspects of male courtship behaviors in wild-type Canton-S males. We found that males take an asymmetric and stereotyped path around a stationary female during courtship whereby they position themselves *∼*1.5 times further away from the female’s head than the tip of her abdomen (*p* < 0.001, 1-Sample T-Test; Figures 1B-C). Next, we trained and used classifiers to identify video frames containing males engaging in three easily-recognizable courtship behaviors: (1) tapping or touching, (2) orienting, and (3) scissoring or wing extensions (Supplemental Figure 1A-E). In our paradigm, these behaviors accounted for *∼*95% of the time that males were actively courting (Supplemental Figure 1F). Additionally, once males began courting, they transitioned between behavioral states at the rates shown in Supplemental Figure 1G.

Spatial analyses of these three courtship elements indicated that bouts of tapping largely occur when the male is on the posterior half of the female, whereas bouts of orienting and scissoring occur when the male is on the anterior half of the female (Figure 1D). Accordingly, tapping positions are largely anti-correlated with scissoring and orienting positions, whereas orienting and scissoring positions are highly correlated with one another (Supplemental Figure 1H-I).

We next sought to determine whether vision was driving the differential spatial positioning of males during specific courtship elements by comparing males courting females under either red light (a condition which limits vision in flies) or white light (Figure 2). We found that under red light, males no longer display an asymmetric courtship path. Instead, they exhibit a symmetric path that is equidistant from both the anterior and posterior ends of the female (*p* < 0.001, 1-Sample T-Test; Figure 2A-C). Further, under red light conditions, the relative angular positions of tapping, orienting, and scissoring behaviors are all uniformly distributed around the female (*p >* 0.05 for all behaviors, Rayleigh Test; Figure 2D). Although red light conditions result in longer latencies to initiate courtship and lower overall courtship indices relative to white light, males still display all three main courtship elements (Supplemental Figure 2A-D). These data indicate that while vision is important for the spatial distribution of courtship elements, it is not required for their release. Nonetheless, when courting under red light, we found that males increase the relative time spent tapping and decrease the time spent scissoring toward the female, while the amount of time spent orienting remained constant (Supplementary Figure 2E). These results are consistent with the observed frequency of transitions from orienting into tapping and orienting into scissoring for males courting under either white or red light (Supplemental Figure 2F-H). Intriguingly, these data suggest that males might compensate for the loss of visual stimuli by increasing chemo- and/or mechanosensory inputs via tapping.

**Figure 2:**
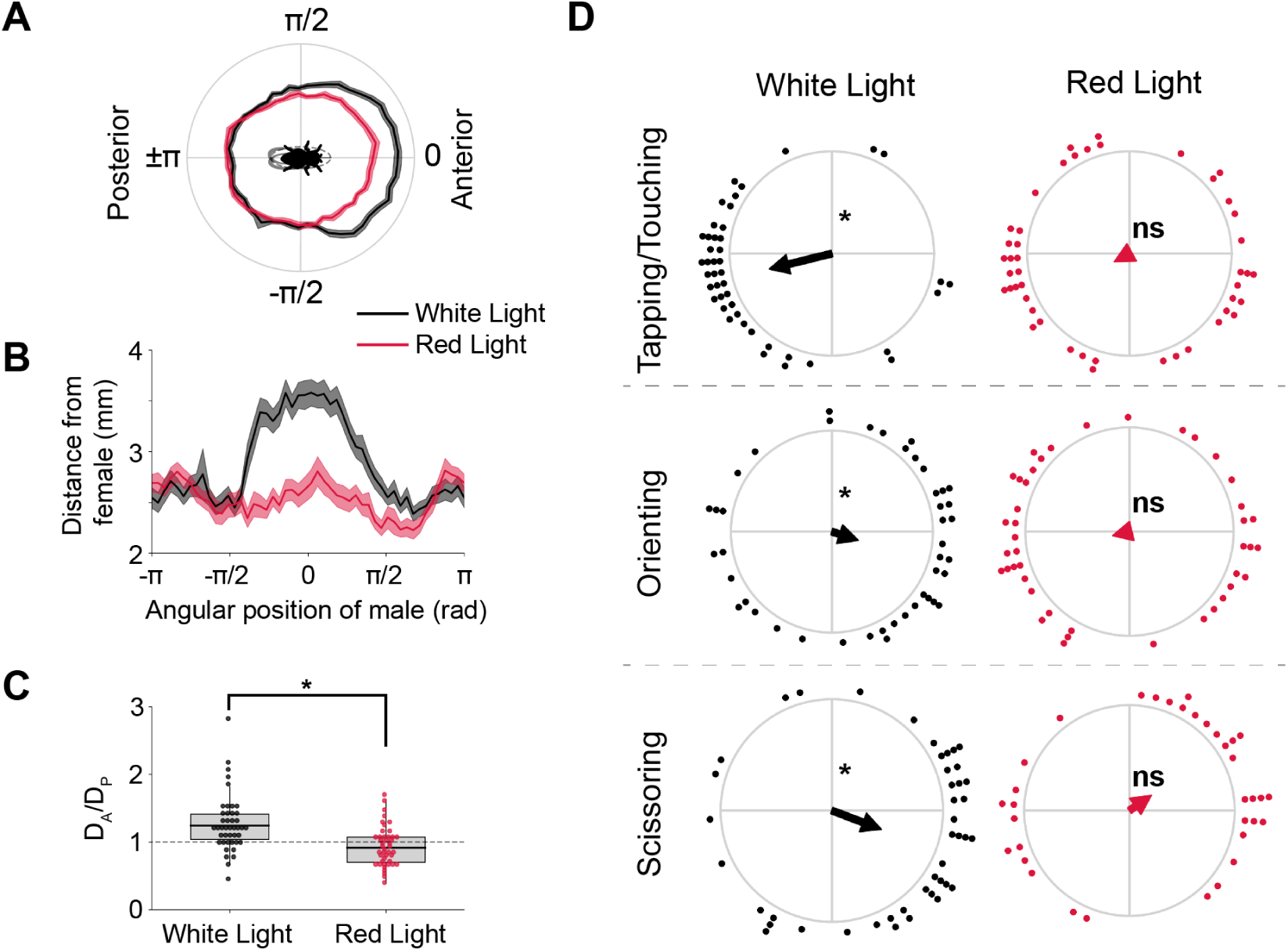
Spatially-stereotyped courtship elements depend on vision. **(A)** Average courtship path for Canton-S males that were allowed to court under either white-(black line) or red-light (red line) (n=48-49/group) **(B)** Same as (A), shown in Cartesian coordinates. **(C)** Maximum distance ratio of male on anterior versus posterior end of female. Males allowed to court under white light had a *D*_*A*_*/D*_*P*_ > 1 (*p* < 0.001, 1-Sample T-Test), whereas males courting under red light had a *D*_*A*_*/D*_*P*_ = 1 (*p* = 0.35, 1-Sample T-Test). These results suggest that males positioned themselves further from the females anterior end only when light is available (*p* < 0.001, One-way ANOVA). **(D)** Average angular positions of males during individual courtship behaviors under either white- or red-light. Under white-light, males positioned themselves to the posterior side of the female during bouts of Tapping/Touching (*p* < 10^-5^, Rayleigh Test), whereas they positioned themselves to the anterior side of the female during bouts of either Orienting or Scissoring (Orienting: *p* < 10^-4^; Scissoring: *p* < 10^-4^; Rayleigh Test). However, males courting under red light did not show directionality in any of their average behavioral locations (Tapping/Touching, Orienting, Scissoring: *p >* 0.1; Rayleigh test). Note that the female is oriented as in (A).

### Males use the eyes of females as a visual marker for directing the spatial distributions of specific courtship elements

Having established that vision plays an important role in the proper spatial positioning of males during courtship, we then asked which specific visual cues direct the precise spatial release of these behaviors. We hypothesized that morphological features on the female body with regions of high contrast could serve as salient visual cues. In *D. melanogaster*, the head contains two dark, red-pigmented eyes that highly contrast with the lighter colored cuticle and could serve as a visual marker for males during courtship (Supplemental Figure 3). We thus tested whether males use the eyes of the females as a visual cue for establishing where to direct specific behavioral elements of the courtship ritual.

First, we asked whether the head of target females is necessary for regulating any spatial aspects of the male courtship ritual (Figure 3A-E and Supplemental Figure 4). We found that when courting headless females under white light, male courtship paths are symmetric and equidistant from either end of the females anterior-posterior axis (Figure 3A-D), and the mean angular positions of tapping, orienting, and scissoring behaviors are uniformly distributed around the female (*p >* 0.05 for all behaviors, Rayleigh Test; Figure 3E). Despite spatial deficits, we found that overall courtship latencies and indices towards intact and headless females are not different (Supplemental Figure 4, A-D). However, we found that males courting headless females exhibit a decreased scissoring frequency, suggesting that anatomical features associated with the head of the female are possibly serving as a visual trigger for the release of this specific courtship element (*p* < 0.05, Kruskal Test; Supplemental Figure 4, E-H).

**Figure 3:**
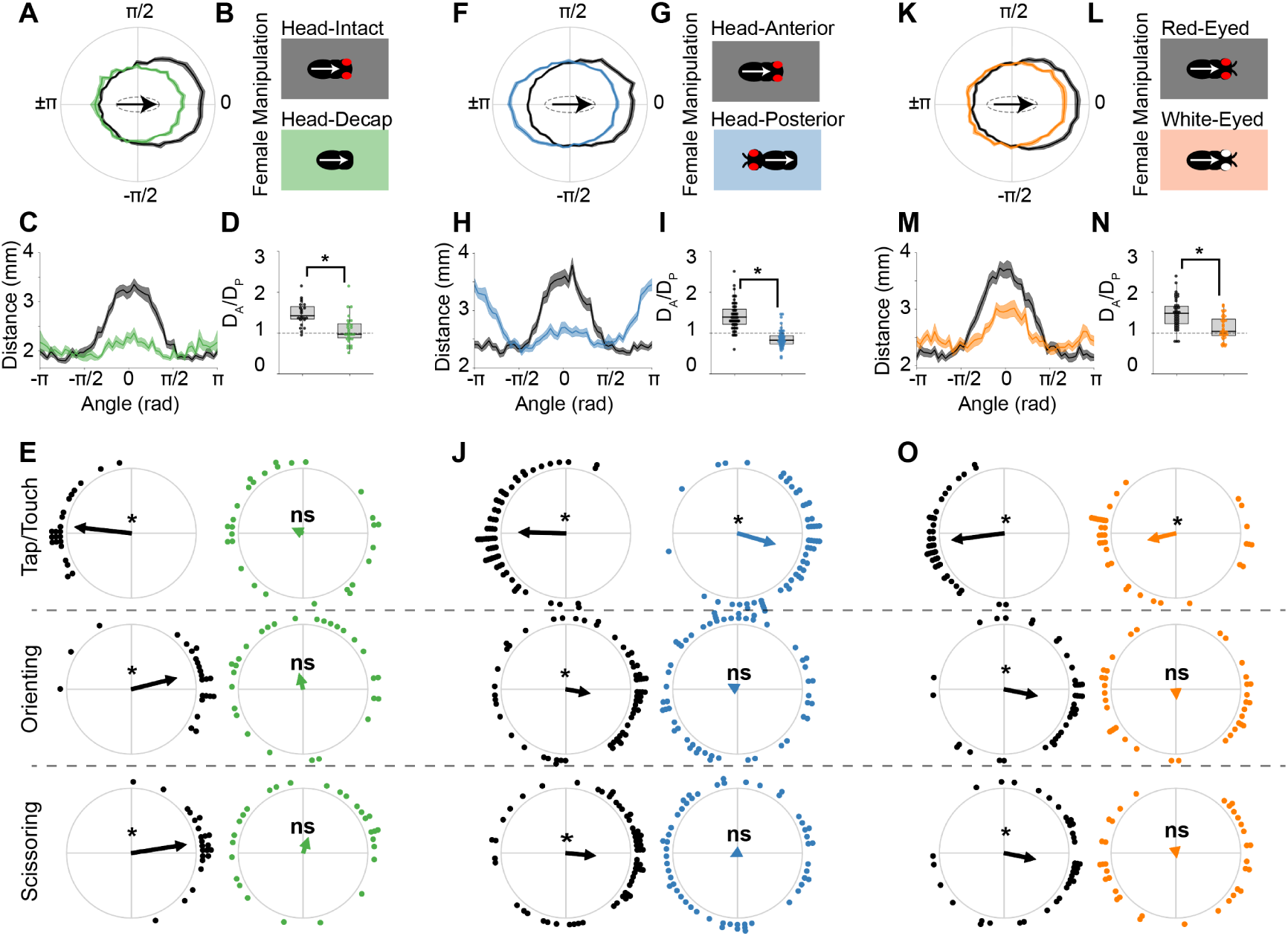
The females head and eye coloration are important visual features for proper male positioning during courtship. **(A)** Average courtship paths of males courting either intact females (Head-Intact, n=32, black line), or females that had been decapitated (Head-Decap, n=32, green line). **(B)** Manipulations for intact and decapitated females. Arrows show the anterior-posterior axis of the female as it has been plotted in (A). **(C)** Average courtship paths of males courting either intact or decapitated females (same as in A), shown in Cartesian coordinates. **(D)** Maximum distance ratio of male on the anterior (*D*_*A*_) versus posterior (*D*_*P*_) end of the female. Males courting intact females positioned themselves significantly further from the females anterior end versus their posterior end (*p* < 10^-5^, 1-Sample T-Test). However, males courting decapitated females did not position themselves significantly further when on either side of the female (*p >* 0.1, 1-Sample T-Test). Thus, males courting intact females had a significantly greater *D*_*A*_*/D*_*P*_ ratio than males courting decapitated females (*p* < 10^-4^, One-way ANOVA). **(E)** Behavioral locations of males courting either intact or decapitated females (asterisks denote significance at *p* < 0.05, Rayleigh Test). Note that females are oriented as in (A). **(F)** Average courtship paths of males courting either intact females (Anterior, n=64, black line), or females that had their heads transplanted to their posterior end (Posterior, n=63, blue line). **(G)** Female manipulations of either the Anterior or Posterior group. Note that arrows are the same as in B. **(H)** Average courtship path of males courting either Anterior or Posterior females (same as in F), shown in Cartesian coordinates. **(I)** *D*_*A*_*/D*_*P*_ ratio for males courting either Head-Anterior (Anterior) or Head-Posterior (Posterior) females. Anterior males positioned themselves significantly further from the females anterior rather than posterior end (*p* < 10^-5^, 1-Sample T-Test), whereas Posterior males positioned themselves significantly more distant from the females posterior rather than anterior end (*p* < 0.01, 1-Sample T-Test). **(J)** Behavioral locations of males courting either Anterior or Posterior females (asterisks same as in E). Note that females are oriented as in (F). **(K)** Average courtship path of males courting either females with red eyes (Red-Eyed, n=47, black line) or white eyes (White-Eyed, n=48, orange line). **(L)** Female manipulations of Red-Eyed and White-Eyed groups. Arrows same as in B. **(M)** Same as in K, shown in Cartesian coordinates. **(N)** *D*_*A*_*/D*_*P*_ ratio for males courting either Red-Eyed or White-Eyed females. Both groups have ratios significantly different from 1 (Red-Eyed, *p* < 10^-5^; White-Eyed, *p* = 0.03; 1-Sample T-Test); however, males in the Red-Eyed group have significantly greater ratios than males in the White-Eyed group (*p* < 10^-5^, One-way ANOVA). **(O)** Behavioral locations of males courting either Red-Eyed or White-Eyed females (asterisks same as in E). Note that females are oriented as in (K).

Next, to determine whether the head of the female is sufficient to induce the release of individual courtship behaviors at their appropriate spatial positions, we examined the behavior of males courting females whose head had been transplanted from her anterior to posterior end. We found that in contrast to males that courted intact females (“Head-Anterior”), males that courted females with a posterior head position (“Head-Posterior”) exhibit an asymmetric courtship path that was biased towards greater distances from the posterior, rather than the anterior, end of her body axis (Figure 3F-I). Thus, males utilize the head as a visual marker to delineate the anterior-posterior body axis of the female during courtship. Further, we found that when courting “Head-Posterior” females, males switch their tapping location to the anterior end of the female body axis. In contrast, the spatial distributions of orienting or scissoring behaviors towards “Head-Posterior” females are randomly distributed (Tapping: *p* < 0.05, Rayleigh Test; Orienting and Scissoring: *p >* 0.05, Rayleigh Test; Figure 3J). These results indicate that the location of the female head is required and sufficient to drive the spatial release of tapping behaviors during courtship. However, female head location is required, but it is not sufficient, to drive the appropriate spatial release of orienting or scissoring behaviors. Despite the significant impact of head location on the spatial organization of male courtship behaviors, there were no significant differences in the overall levels of courtship or individual behavioral elements between groups, suggesting that the presence of the head, independent of its location, is sufficient for maintaining the overall courtship drive of males (Supplemental Figure 5).

Subsequently, we asked which specific visual features of the female head were important for directing the spatial organization of male courtship. Because one of the most striking visual features of the fly head is the contrast between the red-pigmented eyes and the surrounding cuticle (Supplemental Figure 3), we hypothesized that males recognize female eyes during courtship and use this as a visual landmark to coordinate spatial displays during courtship. To test this hypothesis, we generated two congenic lines of wild-type flies that differed in a single mutation in the *white* gene, resulting in red- and white-eyed females with inverted visual contrasts made between the eyes and surrounding cuticle (Supplemental Figure 3). We found that although both female genotypes elicit asymmetric courtship paths, the distances of males from the anterior end of white-eyed females is significantly reduced when compared to males courting red-eyed females (Figure 3K-N). Further, while males courting white-eyed flies tap females at the posterior end, the mean angular locations of both orienting and scissoring bouts were randomly distributed (Figure 3O). In addition, while the relative durations of tapping and orienting were not affected by female eye color, males courting white-eyed females spent significantly less time scissoring during the courtship ritual (*p* < 0.01, Kruskal Test; Supplemental Figure 6). These data are similar to those observed when males courted headless females and suggest that males use the eyes of the female as an important visual cue for coordinating both spatial and temporal aspects of the courtship ritual.

### LC neurons are necessary for regulating spatiotemporal aspects of the male courtship ritual

Recent studies have implicated several classes of Lobula Columnar (LC) visual projection neurons in the regulation of various visually-induced behaviors in the fly [10, 11], including motion detection during specific aspects of courtship [12]. Consequently, we investigated whether LC neurons are also important for the regulation of different spatial and/or temporal aspects of the courtship ritual. Based on previously published associations between the activation of specific LC neurons and behavior, we chose to focus our investigation on five classes of LC neurons that could be involved in regulating various courtship behaviors, including: leg reaching (LC10), forward walking (LC17), backward walking (LC9, LC10, LC16, LC17), turning (LC16, LC17), and looming detection (LC4) [10].

We found that synaptic silencing of specific populations of LC neurons by using targeted transgenic expression of Tetanus Toxin (TNT) [19] led to distinct courtship deficits across all lines we examined. Specifically, we found that different LC neuron subtypes play various roles in mediating both spatial and temporal aspects of the courtship ritual, including regulating the latency to court, fractions of time spent exhibiting each courtship element, and the transition frequencies between each behavioral state (Figure 4 and Supplemental Figure 7). These data suggest that each subpopulation of visual descending neurons is involved in coordinating specific visually-guided behavioral elements of the male courtship ritual.

**Figure 4:**
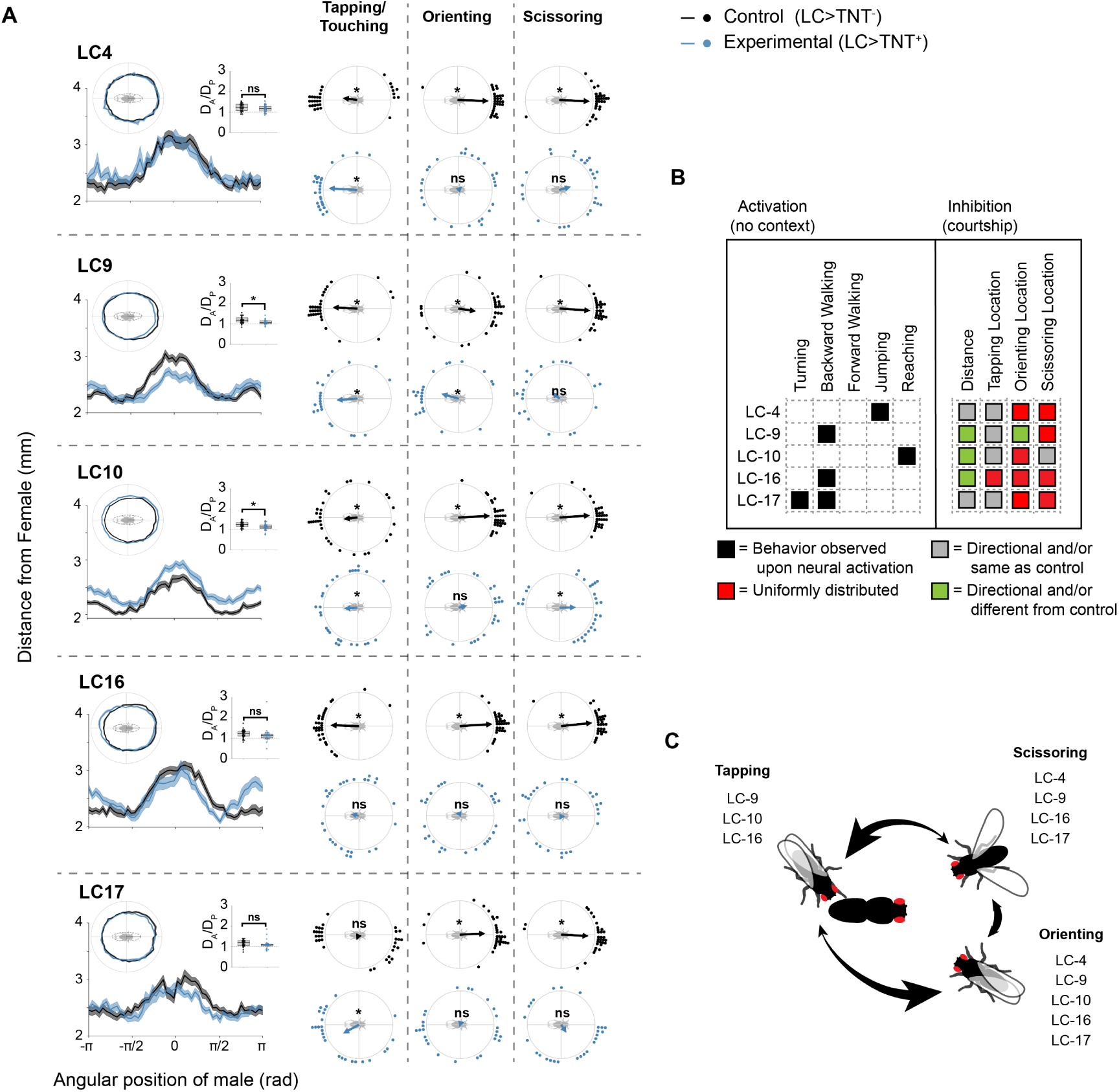
Visual projection neurons are important for directing the spatial localization of male courtship behaviors. **(A)** Average courtship paths, ratios of the maximum male-to-female distance when a male is on either the anterior or posterior end of the female (*D*_*A*_*/D*_*P*_), and average angular positions of the male during Tapping/Touching, Orienting, and Scissoring are shown for each LC line expressing either an inactive (Control, n=32 per line) or active (Experimental, n=32 per line) version of the Tetanus toxin gene (TNT). Asterisks on *D*_*A*_*/D*_*P*_ plots represent significant differences between Control and Experimental groups (*p* < 0.05, One-way ANOVA). Asterisks on circular plots highlight distributions that were significantly different from uniformity (*p* < 0.05, Rayleigh Test). **(B**, left) Solid black squares denote LC lines that engaged in the specified behavior upon neural activation in the absence of a specific behavioral context (data from [10]). **(B**, right) Gray and colored squares denote whether an LC line differed from controls in their spatial positioning during courtship following neural inactivation (data from this paper). **(C)** Summary plot showing which LC lines were important for the proper spatial positioning of males during Tapping, Orienting, and Scissoring.

### LC neurons contribute to stereotyped courtship-related locomotor patterns

The activation or inhibition of LC neurons can elicit or prevent specific movements in flies, respectively [10, 11]. To determine the types of movements required for proper male positioning during courtship, we therefore looked at average movement speeds when males were located at different angular positions around the female (Figure 5A). We found that control males had very stereotypical movement speeds around the female, where they accelerate and achieve high velocities while on either side of the females medial-lateral axis (*V*_*ML*_, areas shown in green in Figure 5B); an effect which is mostly mediated via sideways movement (Figure 5A-C). These males then slow down to much lower velocities when on either the females anterior (*V*_*A*_) or posterior (*V*_*P*_) sides. In contrast, males with inactive populations of LC neurons move slower than controls overall, and while they are capable of speeding up when alongside the females medial-lateral axis (with the exception of LC16), they fail to slow down to the same extent as controls when near the female’s head (as shown by decreased *V*_*ML*_/*V*_*A*_ ratios; Figure 5D). Interestingly, males courting decapitated females show similar movement phenotypes (Figure 5E-G). These results suggest that LC neurons may be responsible for leading to decreased motor output upon detection of the head of the female.

**Figure 5:**
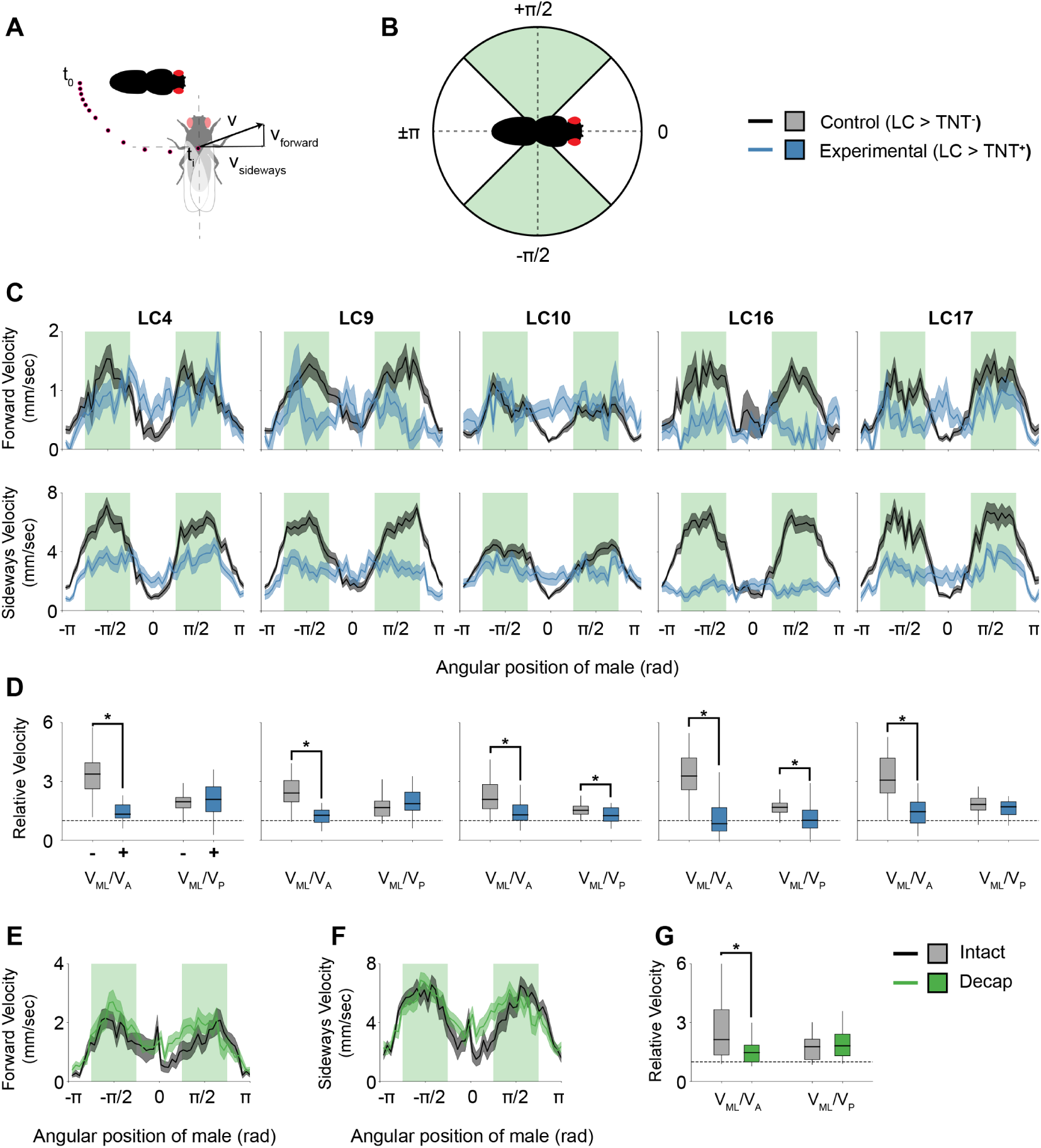
Visual projection neurons mediate movements during courtship. **(A)** Schematic showing the breakdown of the males velocity vector (*v*) at a specific time point (*t*_*i*_) into forward (*v*_*forward*_) and sideways (*v*_*sideways*_) components. **(B)** Schematic highlighting the four spatial quadrants surrounding the female. **(C)** Line plots showing the average velocity (±SEM) of each population of males at each angular bin surrounding the female. Areas of green correspond to the spatial quadrants on either side of the female along the medial-lateral axis; areas of white correspond to spatial quadrants on either side of the anterior-posterior axis. Forward and sideways velocities are affected following the inactivation of most LC lines; however, the most severe deficits occur when males are along either side of the female along the medial-lateral axis. **(D)** Relative velocities showing both (1) the mean sideways velocity when the male is on either side of the females medial-lateral axis (*V*_*ML*_) to the sideways mean velocity when the male is within the females anterior quadrant (*V*_*A*_) and (2) the sideways mean velocity when the male is on either side of the females medial-lateral axis to the mean sideways velocity when the male is within the females posterior quadrant (*V*_*P*_). Inactivation of all lines leads to decreased *V*_*ML*_*/V*_*A*_ratios when compared to controls (*p* < 0.001, One-way ANOVA). **(E-F)** Similar plots to those shown in (C) but for males courting females that were either intact or decapitated (see Figure 3). **(G)** Males courting decapitated females show a significantly smaller *V*_*ML*_*/V*_*A*_ than controls (*p* < 0.05, One-way ANOVA).

## Discussion

Innate behaviors, such as the courtship ritual in *Drosophila melanogaster*, provide unique opportunities to study the sensory cues and neuronal circuits that drive specific behaviors. Here we have identified the contribution of visual cues to the regulation of both spatial and temporal components of the male courtship ritual. Further, we demonstrate that specific classes of visual projection neurons play distinct roles in coordinating the unique spatial patterns of individual behavioral elements of the courtship ritual, as well as the temporal transitions between them. Thus, the studies we present here demonstrate that simple visual cues, such as eye pigmentation, are salient enough signals to direct multiple aspects of a complex social behavior.

Although we primarily focused on the role of the female’s eyes in guiding spatiotemporal aspects of the male courtship ritual, our data also suggest that males must be capable of visually recognizing additional anatomical features of the female body. Specifically, we found that transplantation of the female’s head from the anterior to posterior tip of her body did not lead to the complete reversal of male spatial courtship patterns (Figure 3F-J). Similarly, males were still able to recognize some aspects of the anterior-posterior axis of white-eyed females (Figure 3K-O). One additional anatomical feature on the female that could visually guide males during courtship is the cuticular banding pattern on the abdomen. Although we did not directly test the role of the abdomen in regulating spatial aspects of male courtship, previous studies have indicated that the evolution of pigmentation patterns is under strong sexual selection [20, 21]. Nonetheless, our results suggest that the interplay between pigmentation patterns on both the abdomen and eyes might be utilized by males to direct spatial aspects of their mating displays.

We found that visual cues present on the female are important not only for directing males to specific locations around the female, but they also determine temporal aspects of the courtship ritual. In particular, in the absence of visual input, males decrease the relative amount of time spent engaging in scissoring behaviors and increase the time spent tapping. While these results are most strikingly observed between males courting under either white or red light (Figure 2 and Supplemental Figure 2), we also found that males courting either decapitated or white-eyed females displayed lower levels of scissoring (Figure 3 and Supplemental Figures 4 and 6). One possible interpretation of these results is that males compensate for the loss of sensory information in one modality by collecting more information through the remaining modalities [22]. Therefore, males lacking visual inputs increase their levels of tapping to increase tactile and/or chemosensory inputs from their female courtship target.

Our study suggests that males use a relatively simple visual cue to define the anterior-posterior body axis of courtship targets. Yet the precise spatial and temporal release of specific courtship elements in response to this cue seem to depend on the complex activities of multiple independent populations of LC neurons, each with its own selective effects on male courtship behaviors. While we currently do not know how these circuits interact to generate specific behavioral outputs in the context of courtship, it is likely that each population of visual projection neurons responds to a single visual feature, and the collective activity of these cells is integrated with other contextual cues to generate specific behavioral responses. For instance, as a male approaches a stationary female during courtship, she would appear to be looming. LC4 and LC16 have previously been shown to respond to looming stimuli to generate escape responses [23, 10, 11]; here, we show that in the context of mating, males can also utilize these cell populations to drive distinct courtship behaviors. In contrast to previous studies that have focused on the role of female-generated movement in mediating male chase behaviors [6, 7, 8, 9, 12], our experiments indicate that visual cues play a much broader role in regulating spatial and temporal aspects of mating-related social interactions.

In conclusion, our studies indicate that although vision is not required for triggering male courtship in *Drosophila melanogaster*, it plays an essential role in driving important spatial and temporal aspects of an innate social interaction. Under natural conditions, these previously under-appreciated features of the courtship ritual are likely to be essential for the reproductive success of males. Furthermore, the ability of males to modify the rates at which they engage in particular courtship elements, based on the currently available sensory information, suggest that the plasticity in choosing specific courtship elements may have evolved to enable males to achieve reproductive success even when environmental conditions are not optimal.

## Acknowledgements

We thank the Bloomington *Drosophila* Stock Center (NIH P40OD018537) and the TRiP at Harvard Medical School (NIH/NIGMS R01-GM084947) for providing transgenic fly stocks, and the *Drosophila* Genome Resource Center (NIH grant 2P40OD010949) for plasmids used in these studies. This work was supported by a Howard A. Schneiderman Graduate Fellowship from Washington University to RMM, and NIH grants NS089834 and ES025991 and NSF grants 1545778 and 1707221 to YB-S.

## Author Contributions

RMM designed and conducted the experiments, analyzed the data, and wrote the paper. YB-S designed the experiments and wrote the paper.

## Methods

### Flies

Animals were housed at 25 *o*C and 70% humidity under a 12h:12h light:dark cycle, and reared on a corn-meal based food (Archon Scientific). All flies used in this study are available from Bloomington *Drosophila* Stock Center (BDSC) (see Supplemental Table 1).

Canton-S male and female flies were used for courtship experiments under white and red light (Figures 1-2) and for decapitation and head-transplantation experiments (Figure 3A-J). White-eyed females were derived from *w*^1118^ flies that had been back-crossed into Canton-S for at least 6 generations, and these flies, along with their red-eyed Canton-S counterparts, were used as courtship targets in the red-versus white-eyed experiments (Figure 3K-O). Canton-S females were used as courtship targets in the LC-inactivation experiments (Figures 4-5), and males were derived from crosses between each LC-GAL4 line (LC4, BL68259; LC9, BL68342; LC10-1, BL68337; LC16, BL68331; LC17, BL68356) and either an active (UAS-TNT+, BL28996) or inactive (UAS-TNT-, BL28841) mutant allele of the Tetanus Toxin gene [10, 19].

### Courtship Assay

All courtship trials were conducted at Zeitgeber Time (ZT) 1–5, using 4–6 day old virgin male and female flies. Both males and females were collected immediately following eclosion and moved into 25 mL plastic vials containing corn-meal-based food. Both males and females were kept in single-sex groups of 10–12 for two days, at which time individual males were moved into 5 mL glass vials containing a small amount of fly food and isolated for at least two additional days before testing. On test day, legs and wings were surgically removed from each female target, which was subsequently adhered to a rectangular piece of plastic weigh-boat (approx. 30mm × 30mm) using UV-hardening glue (RapidFix). A circular courtship arena (approx. 23mm diameter × 6mm height) was placed over the fixed female, and males were aspirated into the chamber and allowed to freely court the female for 10 minutes. The orientation of the anterior-posterior body axis of each target female was random across trials.

### Video Recordings

Videos were recorded on a Raspberry Pi NoIR camera with a Navitar 8–48 mm lens for 10 minutes at 24 frames per second and were backlit using LEDs. To record under red-light conditions, LEDs were covered with long-pass, red filters (Neewer).

### Tracking and Behavioral Classification

All videos were analyzed on a per-frame basis using custom software that tracks body and wing positions of courting flies and subsequently classifies whether or not a male was engaging in a particular behavior. Three classifiers were created for identifying frames that contained males engaging in bouts of (1) tapping or touching, (2) stationary orienting, and (3) stationary scissoring/wing extensions. For each frame, several features were calculated from tracking data for use in an AdaBoost decision tree classifier (see Supplementary Table 2). These features were selected based on both previous studies and empirical classifier cross-validation which yielded greater accuracies [24, 25, 26].

Classifiers were created for each experiment by hand-scoring a subset of frames from at least 4 videos containing control males and 4 videos containing experimental males. All classifiers had accuracies >95% (see Supplementary Tables 3-4). To further improve classification accuracies, all videos were hand-scored for bouts of courtship, and any positive behavioral classifications falling outside of courtship were discarded.

Each of the individual courtship behaviors we classified were defined to be mutually exclusive of one another. We specified a behavioral hierarchy whereby tapping/touching took the highest precedence, followed by scissoring/wing extensions, and then orienting. If a video frame contained multiple behavioral classifications, we used the behavioral hierarchy to determine which behavior to retain and eliminated all other classifications from that frame. This was done to eliminate the strong overlap between bouts of scissoring and orienting and allowed for us to more easily determine when a fly transitioned from one behavior to another. Further, this let us calculate pertinent spatial correlations between behaviors (see Supplemental Figure 1).

### Behavioral Data Analysis

Prior to each analysis, the orientation of the female in the courtship arena (along with all tracking data) was rotated such that the females anterior-posterior body axis was aligned along the x-axis, with the head centered at 0 radians and facing to the right (as shown in Figure 1A-B).

#### Courtship Path

The courtship path of each male was calculated by dividing the angular space surrounding the female into 50 bins and taking the mean centroid-to-centroid distance between the male and female during bouts of courtship. For some experiments, both control and experimental males attempted to copulate with the female for extended periods of time. While these males were not physically able to copulate since the female was fixed in place, these long durations of minimal movement had a significant effect on the courtship path, and for our purposes, represented bouts of copulation; they were therefore removed from all analyses.

#### Anterior-Posterior Distance Ratio (D_A_/D_P_)

The *D*_*A*_*/D*_*P*_ ratio was calculated as the ratio of the maximum courtship path when the male was on the front half of the female (*-π/*2 *< θ*_*male*_ *< π/*2) to when the male was on the rear half of the female (*θ*_*male*_ *< -π/*2 or *θ*_*male*_ *> π/*2).

#### Angular Locations of Courtship Elements

The mean angular position of the male with respect to the female was calculated across all frames containing positive classifications for each behavior of interest. Rayleigh values (represented as arrow length in circular plots) were calculated for populations of males.

#### Courtship Latency and Index

The courtship index was calculated as the total fraction of time a male spent courting a female from the first occurrence of any courtship element until the end of the 10-minute trial (*t*_*courting*_*/*(*t*_*end-trial*_ *- t*_*start-courtship*_)). The courtship latency was calculated as the time taken until the first occurrence of courtship during the trail.

#### Behavioral Indexes

Behavioral indexes were calculated for each of the classified behavioral states. These indexes represent the fraction of time a male spent in a particular behavioral state with respect to the duration of time the male spent courting (*t*_*in-state*_*/t*_*courting*_).

#### Behavioral Transitions

The frequency of transitioning from one behavior to another was calculated by taking the number of transitions between each behavior over the total number of behavioral transitions (*n*_*b*1*→b*2_*/n*_*total*_, *n*_*b*2*→b*3_*/n*_*total*_, etc.).

All software and scripts used for tracking, classification, and data analysis were written in Python and are freely available at www.github.com/regginold/drosophila-courtship.

## Supplemental Figures

**Supplemental Figure 1:**
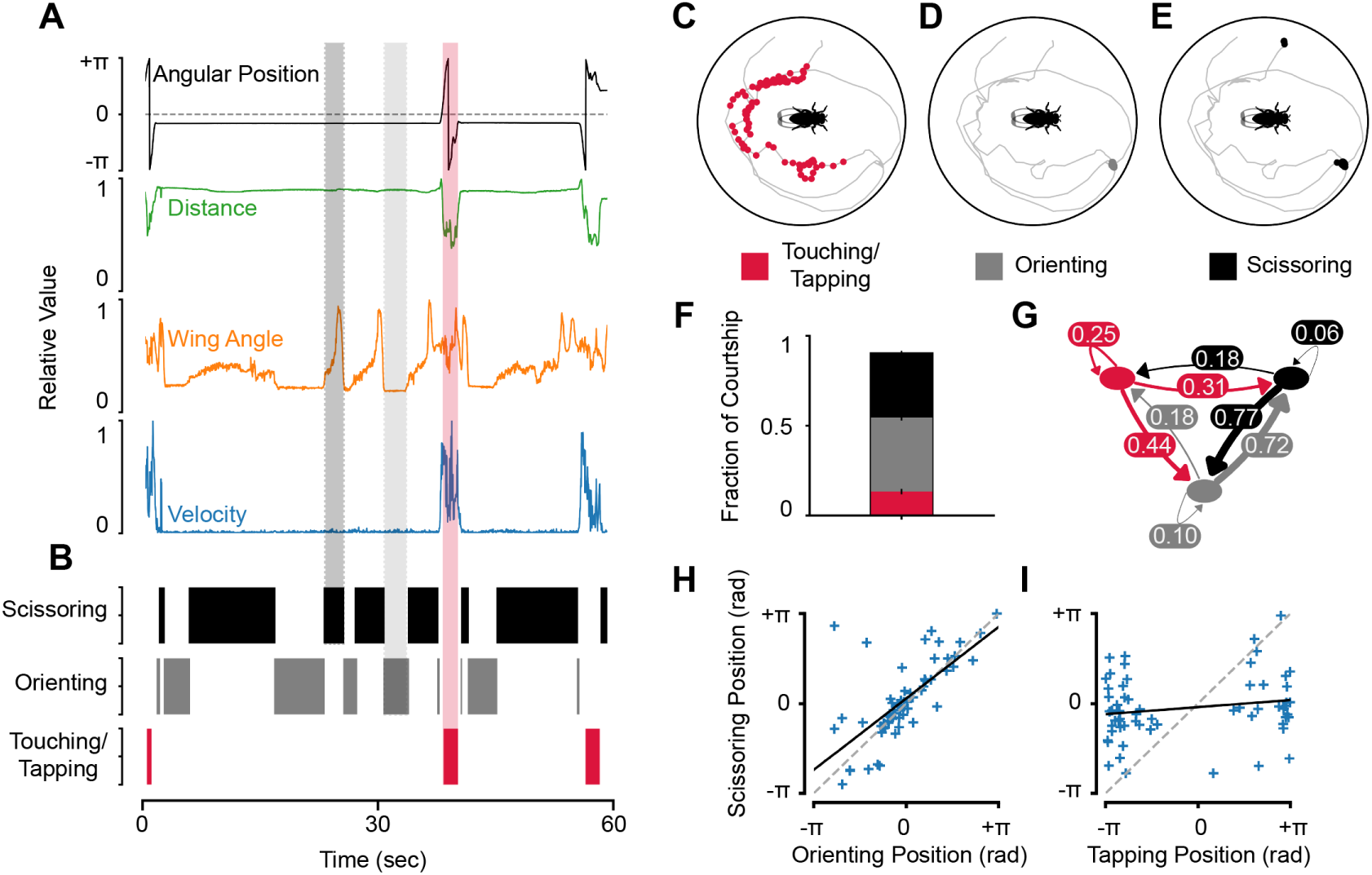
Relationships between individual courtship elements. **(A)** Examples of some of the important features extracted from tracking data. **(B)** Frames classified as Scissoring, Orienting, or Touching/Tapping are shown for an individual male over the course of 1 minute. Behavioral epochs classified as Scissoring are associated with large increases in the angle defined by the males left wing, right wing, and body centroid as well as low velocities. Those classified as Orienting are associated with no changes in the wing angle and low velocities. And those classified as Tapping/Touching are associated with decreased distances between the male and female. **(C-E)** Locations of the male during each of the behavioral epochs from (A-B) are shown along with the track (gray line) produced by the male during the 1 minute segment of courtship. **(F)** Each of the three classified behaviors account for 95% of the behaviors occurring during the courtship ritual. The remaining time is largely spent transitioning between behaviors. **(G)** Transitions between each of the three classified behaviors. **(H)** Mean Scissoring and Orienting positions are highly correlated with one another. **(I)** Mean Scissoring and Tapping positions are highly anti-correlated with one another.

**Supplemental Figure 2:**
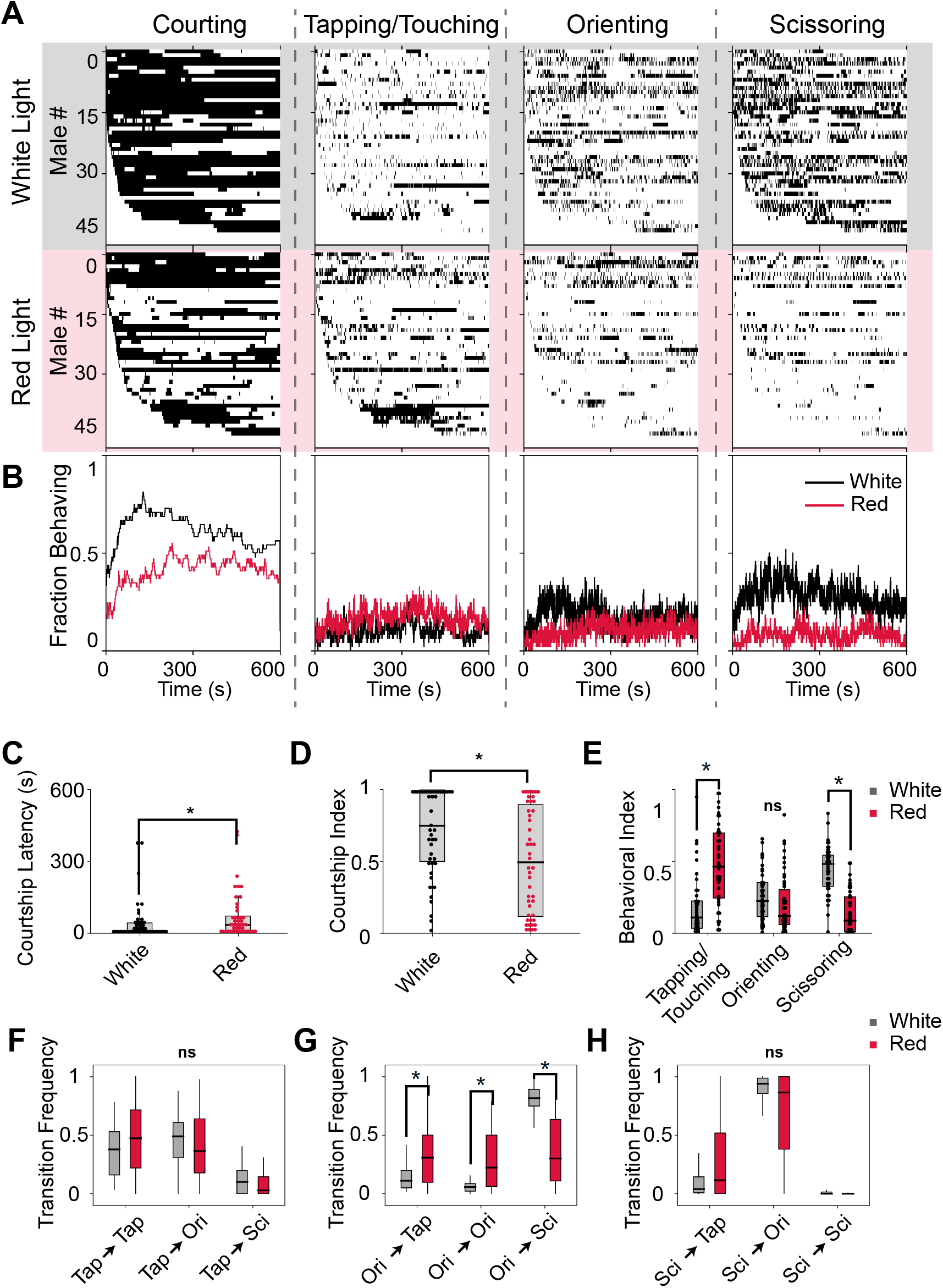
Overall courtship drive and the relative frequencies of individual courtship elements depend on visual inputs. **(A)** Behavioral ethograms are shown for Courtship, Tapping/Touching, Orienting, and Scissoring. Each row represents a courtship trial for one male; areas of black represent frames where the male was engaging in the specified behavior. **(B)** Total fractions of males engaging in each behavior over time. **(C)** Males courting in red light take longer to start courting females (*p* < 0.05, Kruskal Test), and **(D)** court for shorter periods of time (*p* < 0.01, Kruskal Test). **(E)** Behavioral indices are shown for each of the classified behaviors as a fraction of total courtship. Males courting in red light had greater levels of tapping (*p* < 0.001, Kruskal Test) and lower levels of Scissoring (*p* < 0.001, Kruskal Test) than males courting in white light. **(F-H)** Transitions between individual courtship behaviors. Behavioral transitions from Orienting to all other behavioral states are significantly different between males courting in either white or red light (*p* < 0.001, Kruskal Test).

**Supplemental Figure 3:**
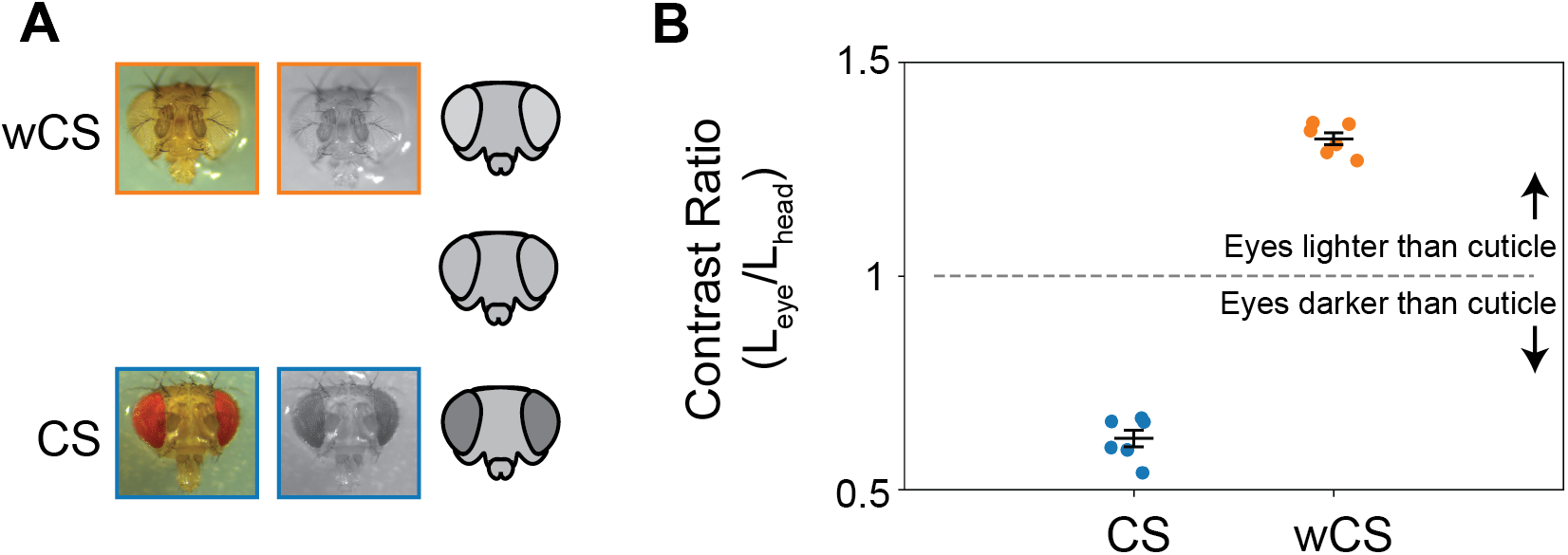
Red-eyed and white-eyed females have inverted eye color contrasts. Canton-S (CS) females were compared to congenic females with a single mutation in the *white* gene (wCS) and Contrast Ratios were calculated for each genotype. **(A)** Images of the heads of CS and wCS females are shown next to a schematic of flies with varying eye contrasts. **(B)** Contrast ratios for CS and wCS flies (n=6/group). Ratios were calculated by converting images to luminances and comparing pixels comprising the eyes (*L*_*eye*_) to pixels comprising the cuticle surrounding the eyes (*L*_*cuticle*_).

**Supplemental Figure 4:**
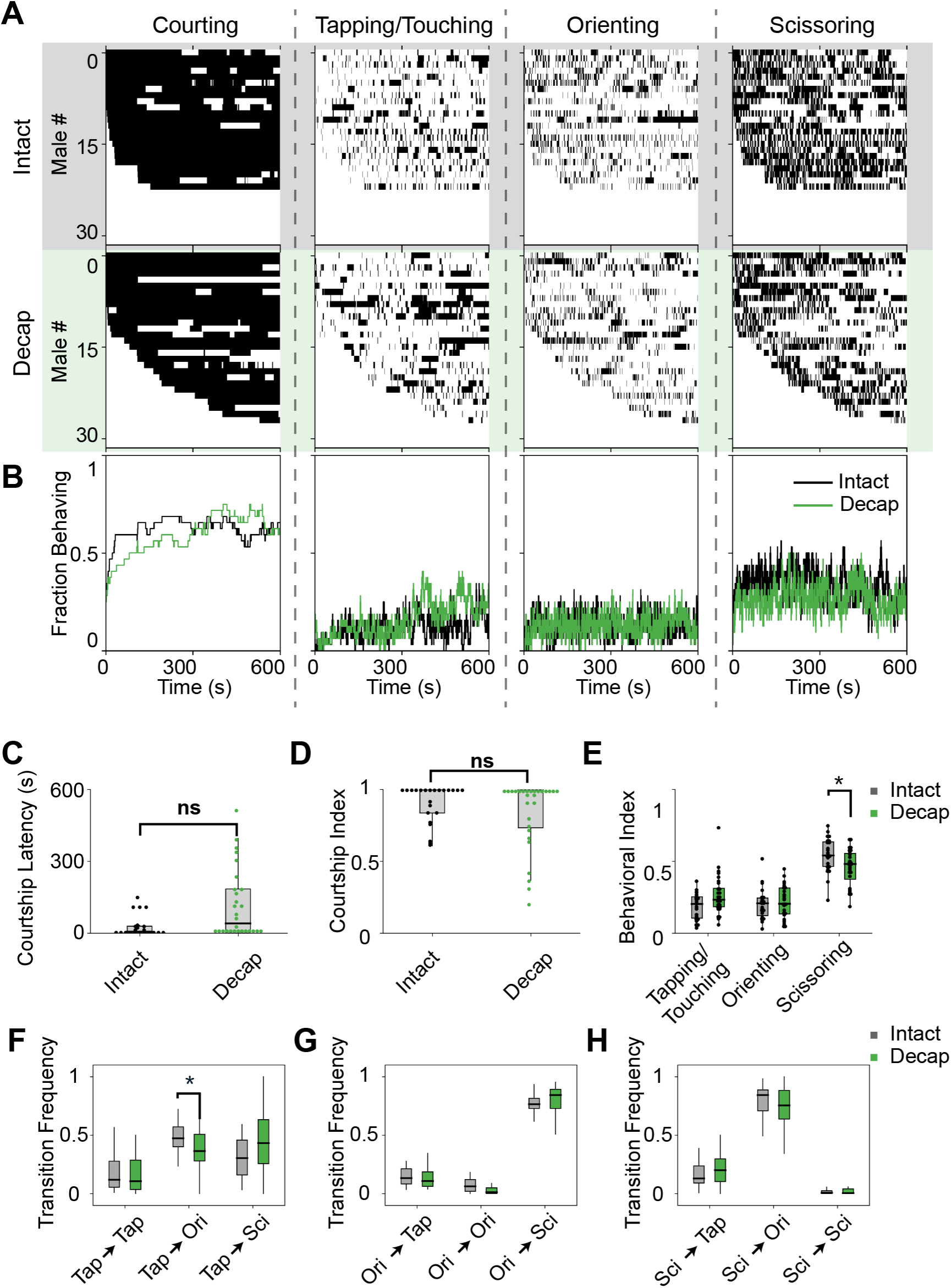
The females head is important for mediating temporal aspects of courtship. **(A)** Behavioral ethograms for males courting either Intact or Decap females. Each row represents an individual fly and each column represents a video frame that has been classified as containing (black) or not-containing (white) a specified behavioral state. **(B)** Fractions of flies engaging in each behavioral state shown in (A) over time. **(C)** Courtship latency and **(D)** courtship index are not significantly different between groups. **(E)** Behavioral indices for touching/tapping and orienting are not significantly different between groups, but males courting decapitated females have lower levels of scissoring (*p* < 0.05, Kruskal Test). **(F-H)** Transitions between individual states of courtship. Males courting decapitated females transition from tapping to orienting less frequently than controls (*p* < 0.05, Kruskal Test), but are otherwise normal.

**Supplemental Figure 5:**
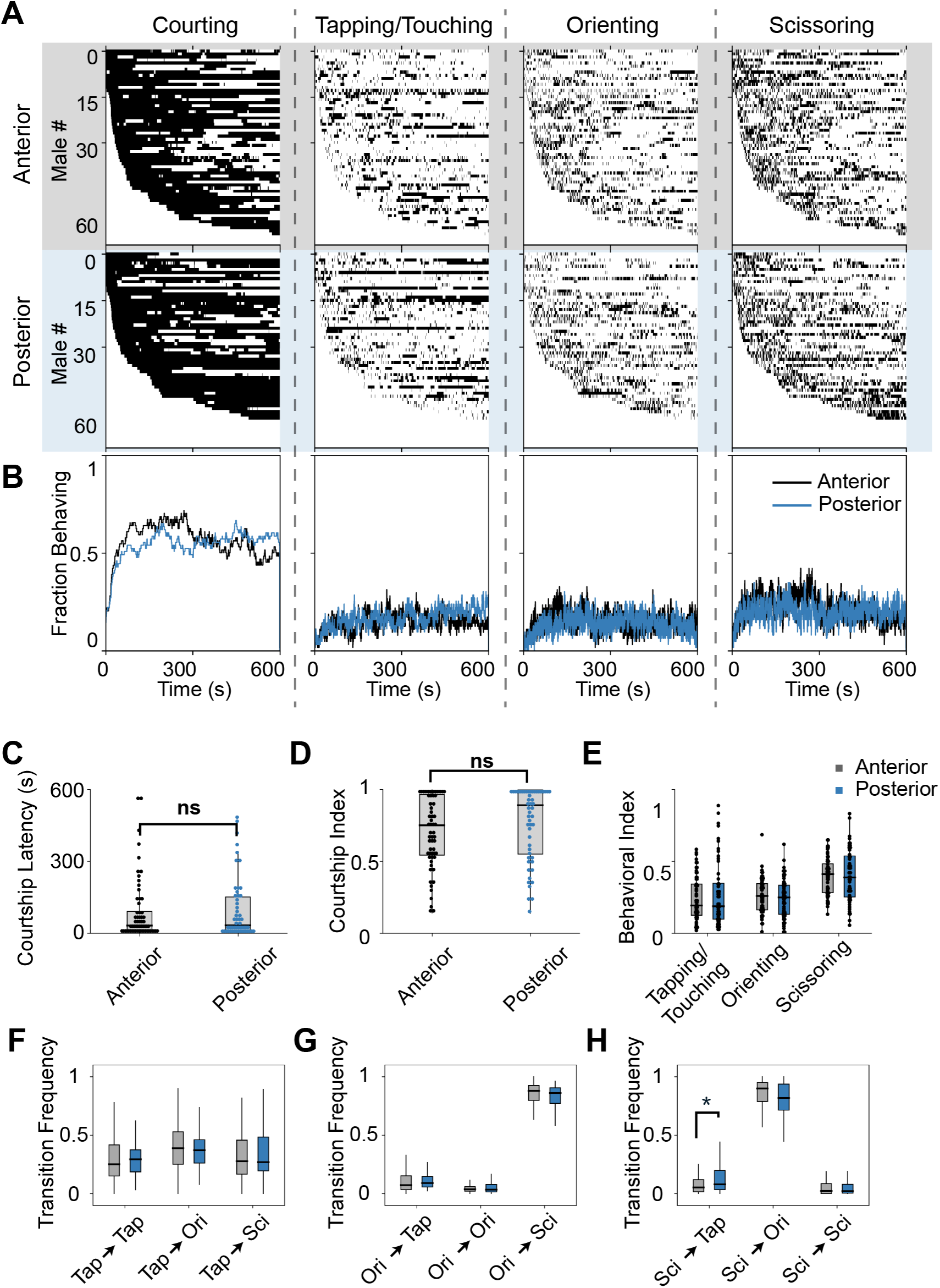
The position of the females head is not required for temporal elements of the courtship ritual. **(A)** Behavioral ethograms for males courting either Head-Anterior (Anterior) or Head-Posterior (Posterior) females. Each row represents an individual fly and each column represents a video frame that has been classified as containing (black) or not-containing (white) a specified behavioral state. **(B)** Fractions of flies engaging in each behavioral state shown in (A) over time. **(C)** The courtship latency, **(D)** courtship index, and **(E)** behavioral indices for tapping/touching, orienting, and scissoring are not significantly different between groups. **(F-H)** Behavioral transition frequencies are shown for each group. Males courting Posterior females show a small but significant increase in transitioning from scissoring to tapping when compared to males courting Anterior females (*p* = 0.047, Kruskal Test).

**Supplemental Figure 6:**
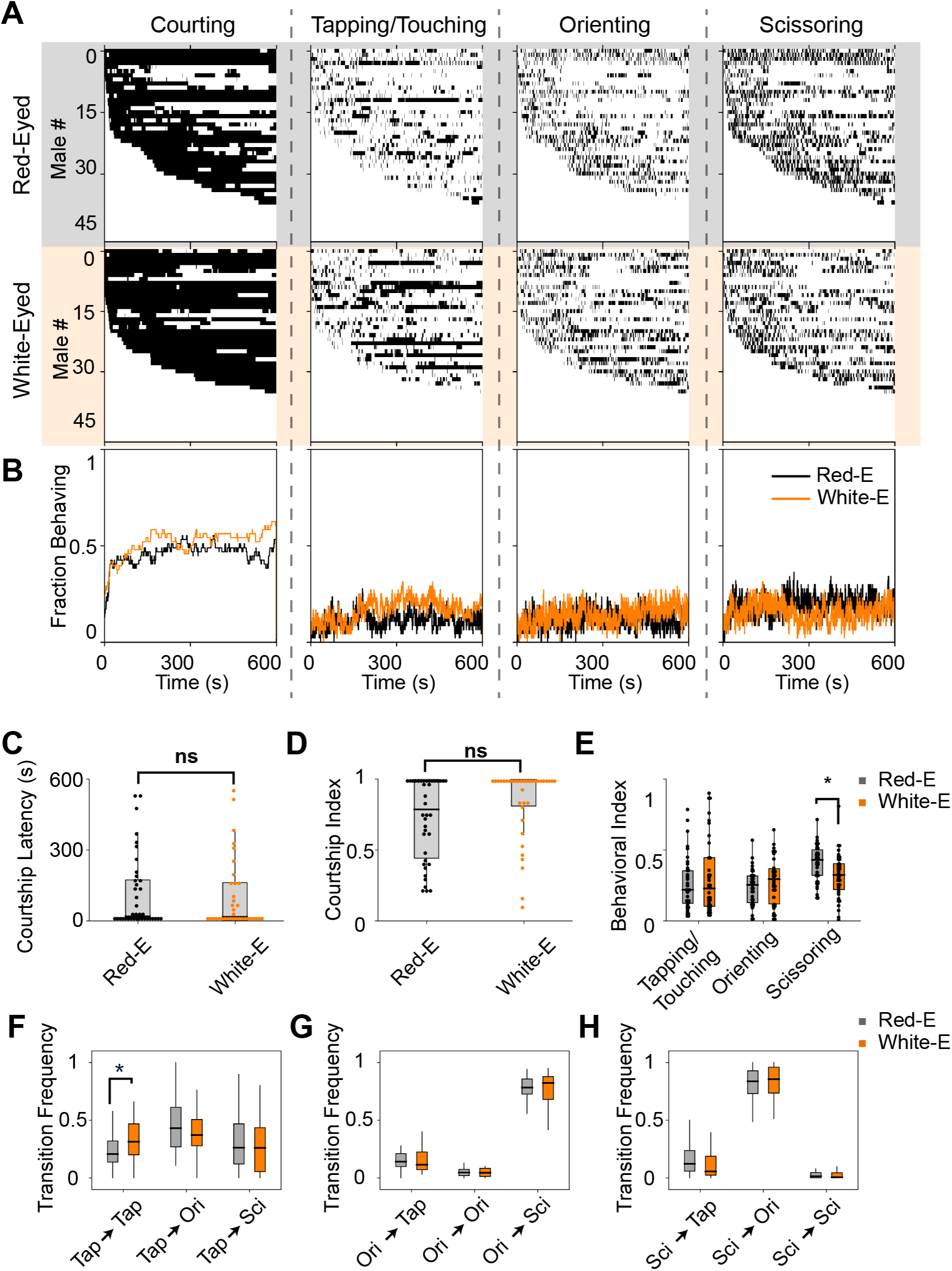
Female eye color is important for mediating temporal aspects of male courtship behavior. **(A)** Behavioral ethograms for males courting either Red-Eyed or White-Eyed females. Each row represents an individual fly and each column represents a video frame that has been classified as containing (black) or not-containing (white) a specified behavioral state. **(B)** Fractions of flies engaging in each behavioral state shown in (A) over time. **(C)** The courtship latency and **(D)** courtship index are not significantly different between groups. **(E)** Males courting White-Eyed females spend less time scissoring compared to controls (*p* < 0.01, Kruskal Test). **(F-H)** Behavioral transition frequencies are shown for each group. Males courting White-Eyed females tend to make more tapping → tapping transitions than males courting Red-Eyed females (shown in F; *p* < 0.05, Kruskal Test).

**Supplemental Figure 7:**
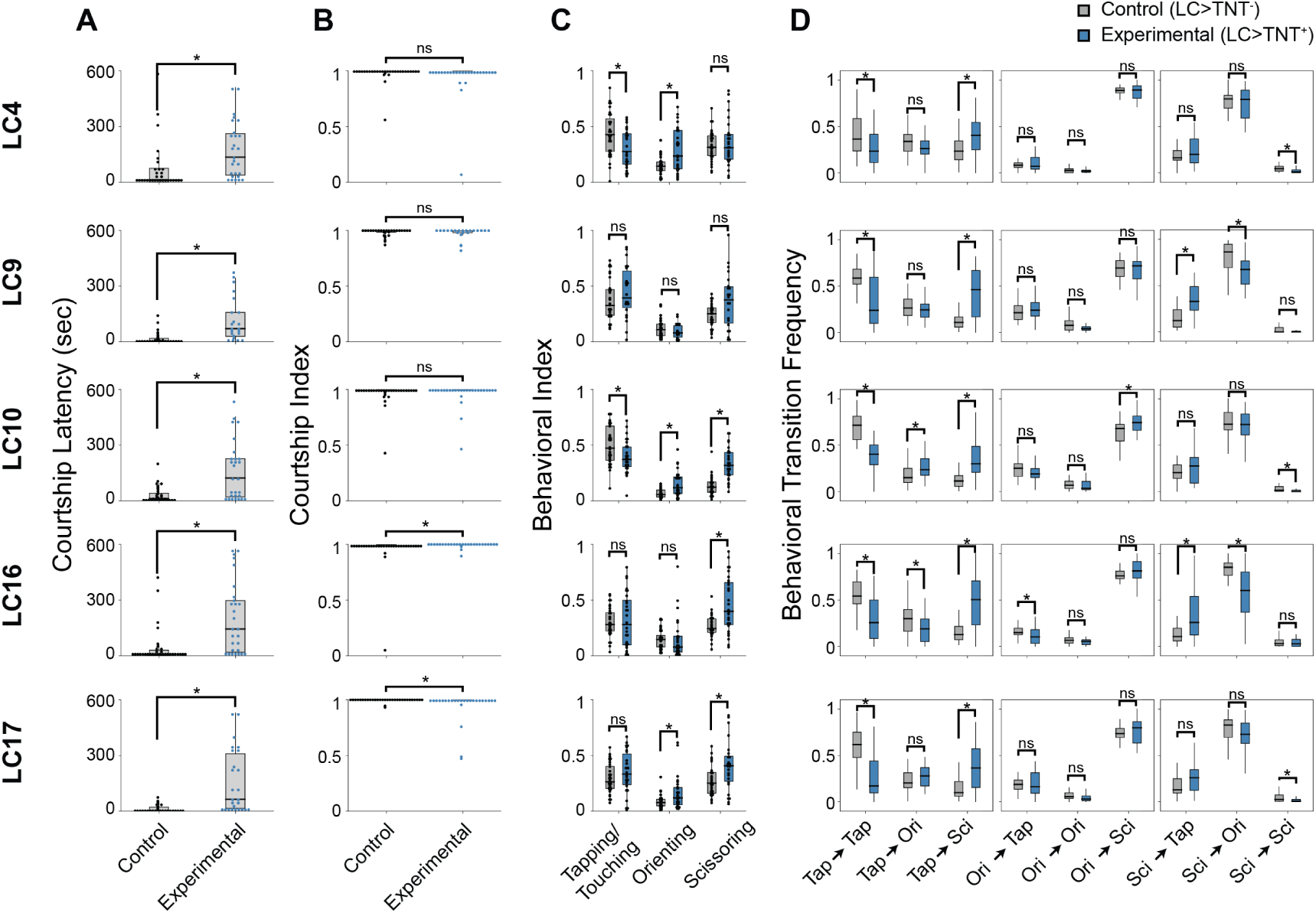
LC neurons are required for temporal aspects of the courtship ritual. **(A)** Inactivation of all LC lines led to significant increases in courtship latencies when compared to controls (*p* < 0.05, Kruskal Test). **(B)** Courtship indices were mostly unaffected by LC inactivation (there were significant differences following inactivation of LC16 (*p* = 0.049) and LC17 (*p* < 0.001); however removal of outliers before statistical testing eliminated this effect). **(C)** Behavioral indices for Tapping/Touching, Orienting, and Scissoring are shown for each LC inactivation. Asterisks represent significant differences between groups (*p* < 0.05, Kruskal Test). **(D)** Frequencies at which flies transitioned between each of the behavioral states during courtship are shown for each LC inactivation.

## Supplementary Tables

**Table 1:** Fly lines used in this paper.

| BDSC # | Description | Reference | Figure(s) |
| --- | --- | --- | --- |
| NA | Canton-S (CS) | NA | 1–5 |
| NA | CS; $w^{1118}$ (wCS) | NA | 3, S3 |
| 68259 | LC4 | Wu et al. [10] | 4–5 |
| 68342 | LC9 | Wu et al. [10] | 4–5 |
| 68337 | LC10-1 | Wu et al. [10] | 4–5 |
| 68331 | LC16 | Wu et al. [10] | 4–5 |
| 68356 | LC17 | Wu et al. [10] | 4–5 |
| 28996 | UAS-TNT<sup>-</sup> | Sweeney et al. [19] | 4–5 |
| 28841 | UAS-TNT<sup>+</sup> | Sweeney et al. [19] | 4–5 |

**Table 2:**
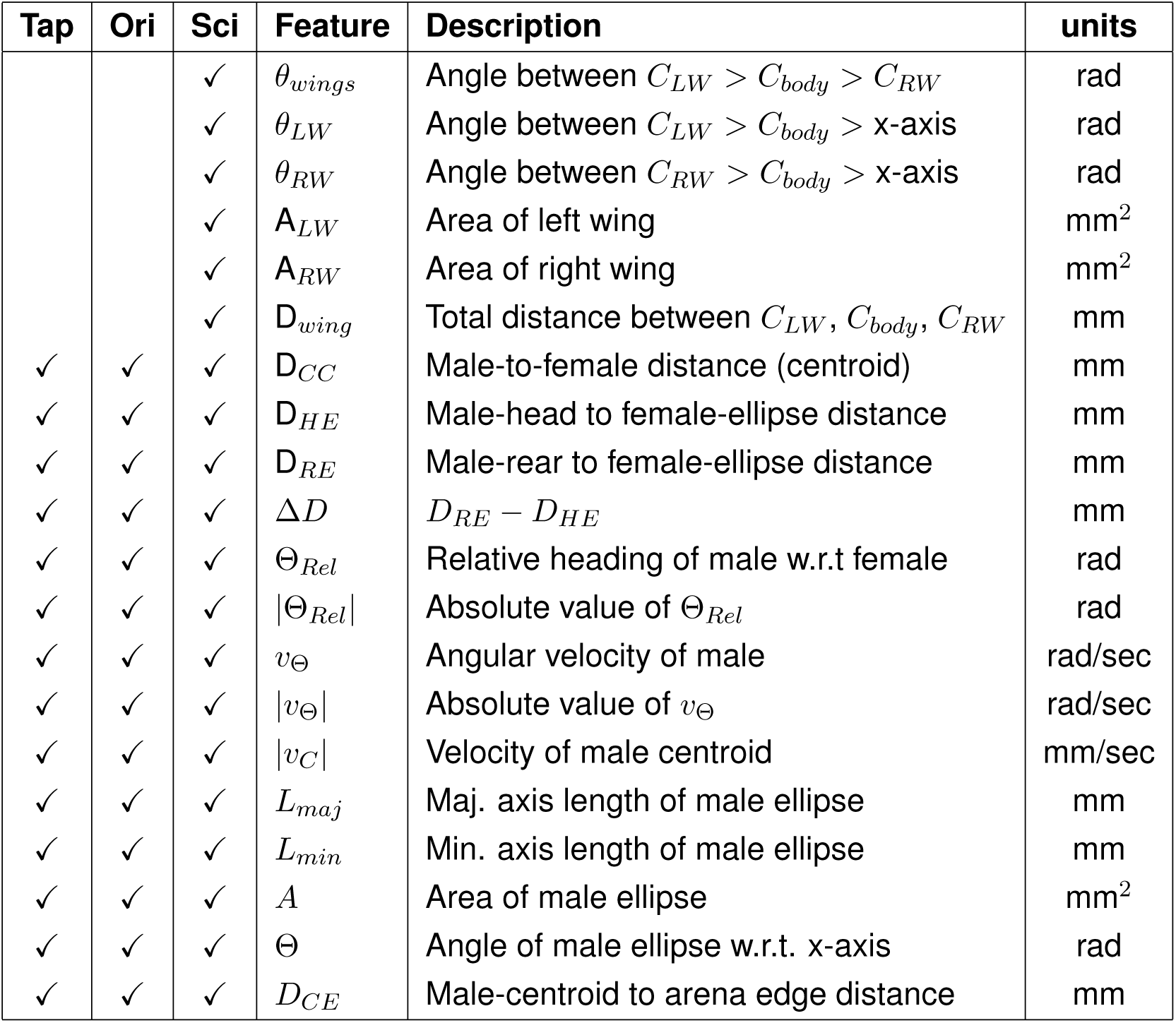
Features used to generate behavioral classifiers. First and second derivatives, as well as windowed statistics, were calculated for all features.

**Table 3:** Behavioral classifier cross-validations. Leave-one-out cross validation was used to determine classifier accuracies, false positive rates (FPRs), and false negative rates (FNRs), as well as standard errors around the mean (*±* SEM) for each of the behavioral classifiers.

| Classifier | $N_{videos}$ Scored | $N_{frames}$ Scored | | Cross Validation | | |
| --- | --- | --- | --- | --- | --- | --- |
|  |  | + | - | Acc (%) | FPR (%) | FNR (%) |
| Tap | 8 | 1137 | 1431 | $95.92 \pm 1.14$ | $2.21 \pm 0.74$ | $1.86 \pm 0.74$ |
| Ori | 8 | 4243 | 4412 | $97.59 \pm 1.57$ | $0.62 \pm 0.58$ | $1.79 \pm 1.55$ |
| Sci | 8 | 4277 | 5248 | $98.18 \pm 1.03$ | $1.37 \pm 0.95$ | $0.46 \pm 0.18$ |

**Table 4:** Behavioral classifier testing. Each classifier was tested on *∼*17,000 hand-scored frames from a single video which was not included in each classifier’s training data set.

| Classifier | Acc (%) | FPR (%) | FNR (%) |
| --- | --- | --- | --- |
| Tap | 99 | 0.763 | 0.211 |
| Ori | 99.4 | 0.128 | 0.507 |
| Sci | 98.9 | 0.696 | 0.434 |

